# TET1 Modulates H4K16 Acetylation by Interacting with hMOF to Regulate Expression of DNA Repair Genes and Oncogenic Transformation

**DOI:** 10.1101/024844

**Authors:** Jianing Zhong, Xianfeng Li, Wanshi Cai, Yan Wang, Shanshan Dong, Jian’an Zhang, Jie Yang, Kangli Wang, Fengbiao Mao, Cheng Zeng, Yuanyuan Li, Jinyu Wu, Huanming Yang, Xingzhi Xu, Zhong Sheng Sun

## Abstract

The Ten Eleven Translocation 1 (TET1) protein is a DNA demethylase that regulates gene expression through alteration of DNA methylation. Recent studies have demonstrated that TET1 could modulate transcriptional expression independent of its DNA demethylation activity; however, the detailed mechanisms underlying TET1’s role in such transcriptional regulation remain not well understood. Here, we uncovered that Tet1 formed a chromatin complex with histone acetyltransferase Mof and scaffold protein Sin3a in mouse embryonic stem cells by integrative genomic analysis using publicly available ChIP-seq data sets. Specifically, the TET1/SIN3A/hMOF complex mediates acetylation of histone H4 at lysine 16, *via* facilitating the binding of hMOF on chromatin, to regulate expression of important DNA repair genes in DNA double strand breaks, including *TP53BP1*, *RAD50*, *RAD51*, and *BRCA1*, for homologous recombination and non-homologous end joining repairs. Under hydrogen peroxide-induced DNA damage, dissociation of TET1 and hMOF from chromatin, concurrent with increased binding of SIRT1 on chromatin, led to hypo-acetylation of H4K16, reduced expression of these DNA repair genes, and DNA repair defects in a DNA methylation independent manner. A similar epigenetic dynamic alteration was also observed in H-RAS^V12^ oncogenic-transformed cells, supporting the notion that suppression of TET1 downregulates DNA repair genes through modifying H4K16ac, instead of its demethylation function, and therefore contribute to tumorigenesis. Taken together, our results suggested a mechanistic link between a novel TET1 complex and H4K16ac, DNA repair genes expression, and genomic instability.

## INTRODUCTION

The Ten Eleven Translocation 1 (TET1) protein, a member of TET family, is a key player in DNA demethylation(Veron and Peters 2011). However, a recent study revealed that Tet1, in addition to its transcriptional regulatory function through its catalytic activity in DNA demethylation, possesses both activation and repressor functions in the regulation of a certain subset of genes in mouse embryonic stem cells (mESCs)(Williams et al. 2011). This observation was further supported by a study in which changes of transcriptional expression induced by overexpression of TET1 were highly similar to those induced by its demethylation-enzymatically-dead mutant in differentiated cell lines, suggesting that TET1 mainly regulates gene expression through a DNA methylation independent manner(Jin et al. 2014). The repressive role of TET1 in transcriptional regulation has been proposed to derive from its interaction with polycomb repressive complex 2 (PRC2) to form a histone modifying complex, thereby modifying chromatin repressive mark (H3K27me3) in mESCs(Wu et al. 2011). However, the interaction between TET1 and PRC2 complex is, so far, exclusively presented in embryonic stem cells (ESCs), but not in differentiated cells such as fibroblasts and HEK293T cells(Neri et al. 2013), indicating that TET1/PRC2 complex may act to repress gene expression in an ESCs-specific manner. On the other hand, SIN3A (homolog of Sin3 in yeast), a key component in multiple regulatory complexes, is involved in both transcriptional repression and activation through recruitment of diverse transcriptional factors or chromatin remodeling machinery at target promoters(Kadamb et al. 2013; Solaimani et al. 2014). A recent study has shown that TET1 interacts with SIN3A in both mESCs and HEK293T cells and presents highly overlapping binding profile on genome-wide(Williams et al. 2011), implying TET1 may associate with SIN3A to regulate gene expression in both ESCs and differentiated cells. However, the exact mechanisms underlying the functional nature of TET1 and its associated protein complexes in regulating its target gene expression remain to be unveiled.

Recently, it was demonstrated that there are dysfunctional DNA repair mechanisms and increased mutation frequencies in TET1-deficient non-Hodgkin B cell lymphoma (B-NHL), indicating that TET1 function as a tumor suppressor(Cimmino et al. 2015). This observation, in line with a previous study in which there were decreased foci of MLH1 and delayed removal of RAD51 in mouse Tet1-knockout primordial germ cells(Yamaguchi et al. 2012), indicates that TET1 plays an important role in DNA repair in mammalian cells. However, the underlying mechanisms of TET1 functions in DNA repair in response to DSBs are unclear.

Homologous recombination repair (HRR) and non-homologous end joining (NHEJ) are two categories of DNA repair pathway in response to DNA double strand breaks (DSBs). Some DNA repair genes, such as *RAD50*, *BRCA1*, *RAD51*, and *TP53BP1*, act as tumor suppressors and are frequently mutated or aberrantly downregulated in human cancers, resulting in impairments of DNA repair in response to DSBs, which is recognized as one of the hallmarks of tumorigenesis (Hanahan and Weinberg. 2011; Negrini et al. 2010). Whole Genome Bisulfate Sequencing (WGBS) data analysis in the Tet1-deficient primordial germ cells showed that the methylation levels of most DNA repair genes had no obvious alteration (Yamaguchi et al. 2012), indicating that Tet1 possibly affects expression of DNA repair genes through a mechanism independent of its DNA demethylation function.

H4K16ac is a well-known targeted epigenetic site for post-translational modification in transcriptional activation(Taylor et al. 2013). Human MOF (hMOF, also known as KAT8), a member of the MYST (Moz-Ybf2/Sas3-Sas2-Tip60) family of HATs, specifically modifies H4K16ac and is frequently downregulated in various types of cancers, including medullo-blastomas, breast carcinomas, colorectal carcinoma, gastric cancer, and renal cell carcinoma(Cao et al. 2014; Pfister et al. 2008). Studies have shown that depletion of hMOF renders both a global reduction of H4K16ac and DNA repair defects in budding yeast and mammal cells (Li et al.; Sharma et al. 2010). In addition, overexpression of hMOF reverses silencing of certain tumor suppressor genes induced by H4K16 deacetylation (Kapoor-Vazirani et al. 2008). Conversely, SIRT1 has the ability to deacetylate H4K16ac (Vaquero et al. 2007;Mishra et al. 2014), and is required for DNA repair and genomic stability in both yeast and mammals(Uhl et al. 2010; Boulton and Jackson 1998). Noteworthy, an elevated SIRT1 expression has been observed in a variety of human cancers relative to their non-transformed counterparts, including leukemia, glioblastoma, prostate, colorectal, and skin cancers(Chen et al. 2005; Huffman et al. 2007; Liu et al. 2009). Importantly, exogenous expression of SIRT1 reverses the effects of hMOF on H4K16ac and sensitization to the topoisomerase II inhibitor of cancer cells(Hajji et al. 2010), implying H4K16ac is dynamically modulated by both hMOF and SIRT1.

In this study, we first revealed, through integrative genomic analysis using publicly available ChIP-seq data sets, that significantly overlapped distribution of TET1, Sin3a, Mof, and H4K16ac were observed in mESCs. We further demonstrated that TET1, hMOF, and SIN3A interact with each other by *in vitro* biochemical studies in human cell lines. We next showed that the identified TET1 chromatin complex specifically modulates H4K16ac. Finally, we uncovered, under DNA damage and oncogenic-induced transformation, that dynamic recomposition of these TET1/SIN3A/hMOF chromatin complex components could cause hypo-acetylation of H4K16, thereby impairing DNA repair and ultimately involving in tumorigenesis in a DNA methylation independent manner.

## RESULTS

### Integrative genomic analysis reveals similar binding patterns between Tet1, Mof, and H4K16ac in mESCs

Previous studies have generated a considerable number of ChIP-seq data sets of DNA binding proteins (DBPs) from mESCs, which is available in GEO and ENCODE databases. We retrieved all 219 available ChIP-seq data sets corresponding to 103 different DBPs in mESCs to investigate the co-occupancy between Tet1 and the rest of the DBPs (**Supplementary Table. 1**). In hierarchical clustering, pair-wise correlation analysis between Tet1 and other DBPs demonstrated that Tet1 could be grouped into one sub-cluster with 13 DBPs in 2000bp transcriptional start site (TSS) flanking region (defined as promoter) (Fig. 1a). Next, the same correlation analysis of each component in this sub-cluster and six available histone modifications in mESCs showed that Tet1, with other five DBPs (Kdm2a, Mof, Dpy30, Sin3a, and Lsd1), were closely related to H4K16ac, H3K4me3, H3K9ac, and H3K27ac (Fig. 1b). Given that Mof is the only histone acetyltransferase in the sub-cluster, whereas Kdm2a, Lsd1, and Dpy30 are either histone demethylases or histone methyltransferase complex component, we decided to focus on the investigation of the co-efficiency among the three histone acetylation marks and Tet1, Mof, and Sin3a by correlation analysis. Our results revealed that Tet1, Sin3a, Mof, and H4K16ac were clustered together with the highest correlation coefficients (Fig. 1c). Furthermore, ChIP-seq signals enrichment analysis revealed that Tet1, Sin3a, Mof, and H4K16ac had highly overlapping distribution patterns around promoter regions (Fig. 1d and **Supplementary Fig. 1**). Taken together, these observations imply that TET1, SIN3A, hMOF have highly similar binding patterns with H4K16ac at the genomic level.

**Figure 1.**
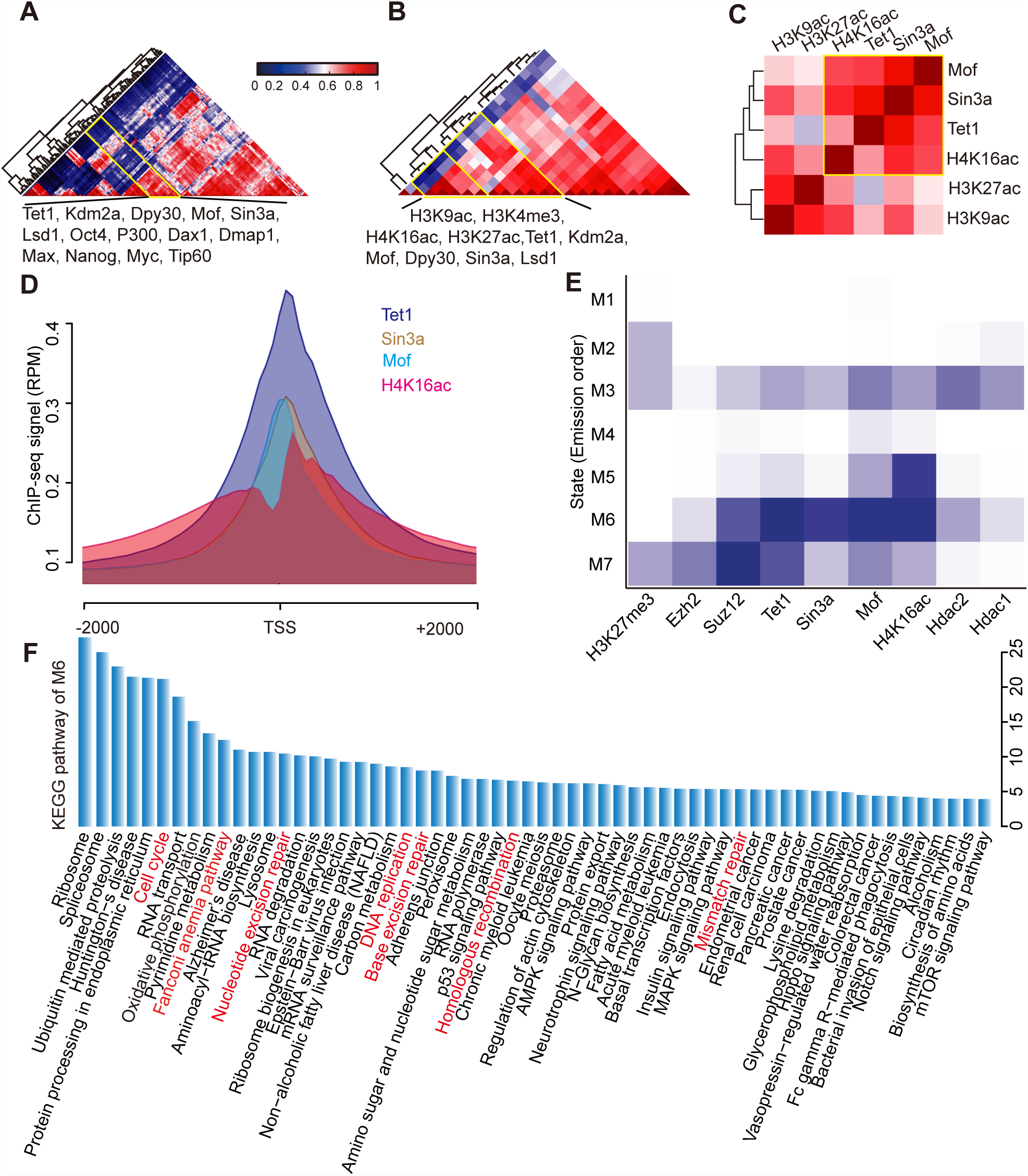
Integrative genomic analysis of published ChIP-seq data sets in mESCs show highly similar binding patterns between Tet1 and H4K16ac. (**a**) Pearson correlation between Tet1 and other 102 DBPs (**Supplementary Table 1**) in 2000bp TSS flanking regions. deepTools was employed to calculate the correlation of ChIP-seq data. Color bar represented correlation coefficient. (**b**) Further correlation between Tet1, other 13 DBPs (Kdm2a, Dpy30, Mof, Sin3a, Lsd1, Oct4, p300, Dax1, Dmap1, Max, Nanog, Myc, and Tip60) and 6 histone modifications (H3K4me3, H3K9me3, H3K9ac, H3K27ac, H3K36me3, and H4K16ac) in 2000bp TSS flanking regions. (**c**) Correlation of Tet1, Sin3a, Mof, H4K16ac, H3K27ac and H3K9ac. (**d**) The bindings of Tet1, Sin3a, Mof, and H4K16ac are commonly enriched in the TSS regions. (**e**) Seven distinct distribution among Tet1/Sin3a/Mof complex, PRC2 complex, and Sin3a/Hdac1/Hdac2 complex were generated by ChromHMM. (**f**) The enriched pathways in cluster M6, including DNA repair pathways marked as red color.

In order to determine whether TET1/SIN3A/hMOF was a complex distinguished from either PRC2 complex or SIN3A/HDAC complex and targeted different chromatin marks, we investigated the binding profiles of the major components of these three complexes, including Suz12, Ezh2, Sin3a, Tet1, Mof, Hdac1, and Hdac2, as well as their associated histone marks H3K27me3 and H4K16ac. Initially, we determined binding states by dividing promoter regions from ChIP-seq data set of each of above proteins and histone marks into clusters M1-M7 using ChromHMM (**Supplementary Fig. 2a, 2b**). Our results showed that Tet1/Sin3a/Mof complex, PRC2 complex, and Sin3a/Hdac1/Hdac2 complex were enriched in cluster M6, M7, and M3, respectively (Fig. 1e). Remarkably, Tet1, Mof, and Sin3a combined with H4K16ac, commonly enriched in cluster M6, was primarily related to DNA damage and repair associated pathways by KEGG pathway analysis (Fig. 1f). In addition, H4K16ac was enriched in M5, which was significantly associated with DNA repair related biological processes similar to those in M6 (**Supplementary Fig. 2c**). Intriguingly, we observed most of promoter regions in 177 DNA repair genes, including *Brca1*, *Brca2*, *Rad51*,*Trp53*, and *Mlh1,* as shown in **Supplementary Fig. 3**, were co-occupied with binding of Tet1, Sin3a, Mof, and H4K16ac. In summary, our integrated genomic analysis indicates that Tet1 may form a complex with Mof and Sin3a targeting H4K16ac in mESCs.

**Figure 2.**
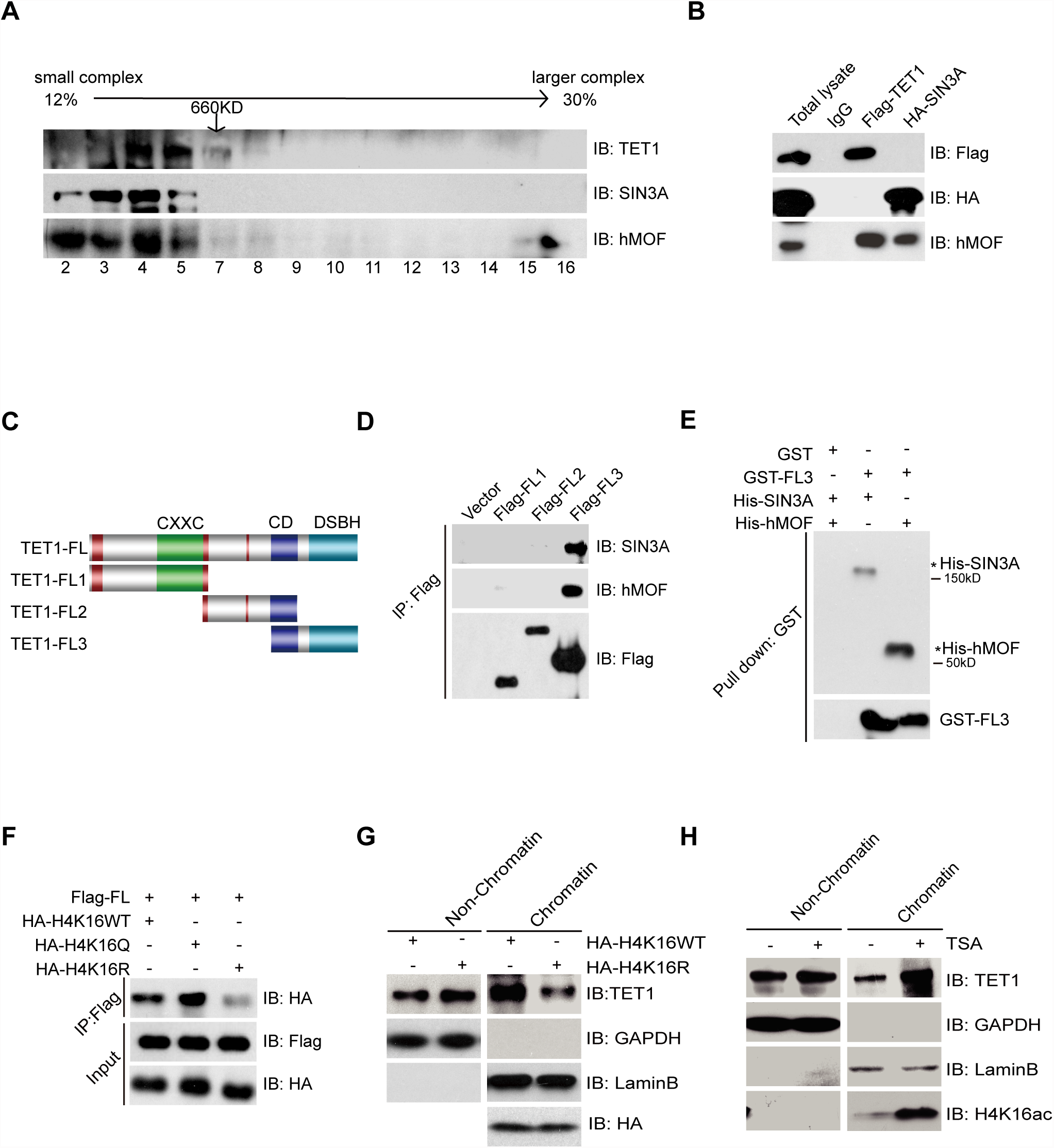
TET1 forms a chromatin complex with hMOF, and SIN3A targeting H4K16ac. (**a**) Nuclear extracts from HEK293T cells were added to a 12%–30% sucrose gradient, and fractions were assayed by immunoblotting. The fraction numbers and 660 kDa molecular mass standard are given across the top. The larger fraction numbers indicate the fraction with smaller molecular weight. (**b**) TET1 and SIN3A were shown interacting with hMOF using nuclear protein immunoprecipitations (IPs) in TET1 overexpressing HEK293T cells. (**c**) Schematic representation of TET1 fragments, including Flag-FL1, Flag-FL2, and Flag-FL3 (CXXC: binding CpG islands; CD: Cysteine-rich domain; DSBH: double stranded β-helix). (**d**) HEK293T cells were transiently transfected with Flag-FL1, Flag-FL2, and Flag-FL3, respectively. Nuclear protein IPs were performed using a Flag-tag antibody, followed by Western blot analysis using indicated antibodies. (**e**) GST and GST-FL3 were expressed in BL21 cells and purified following pGEX-GST-vector’s manual. His-SIN3A and His-hMOF1 were also expressed in BL21 and purified and. Pull down assays were performed using a GST-tag antibody. (**f**) Co-transfections with Flag-TET1-FL and H4K16WT, H4K16Q, or H4K16R were carried out followed by IPs using a Flag-tag antibody. (**g**) Western blot analysis of the fractions in HA-H4K16R and HA-HK16WT overexpressing cells using antibodies as indicated. (**h**) Western blot analysis of the fractions before and after treatment with 1μM TSA for 1h in HEK293T cells using antibodies as indicated.

**Figure 3.**
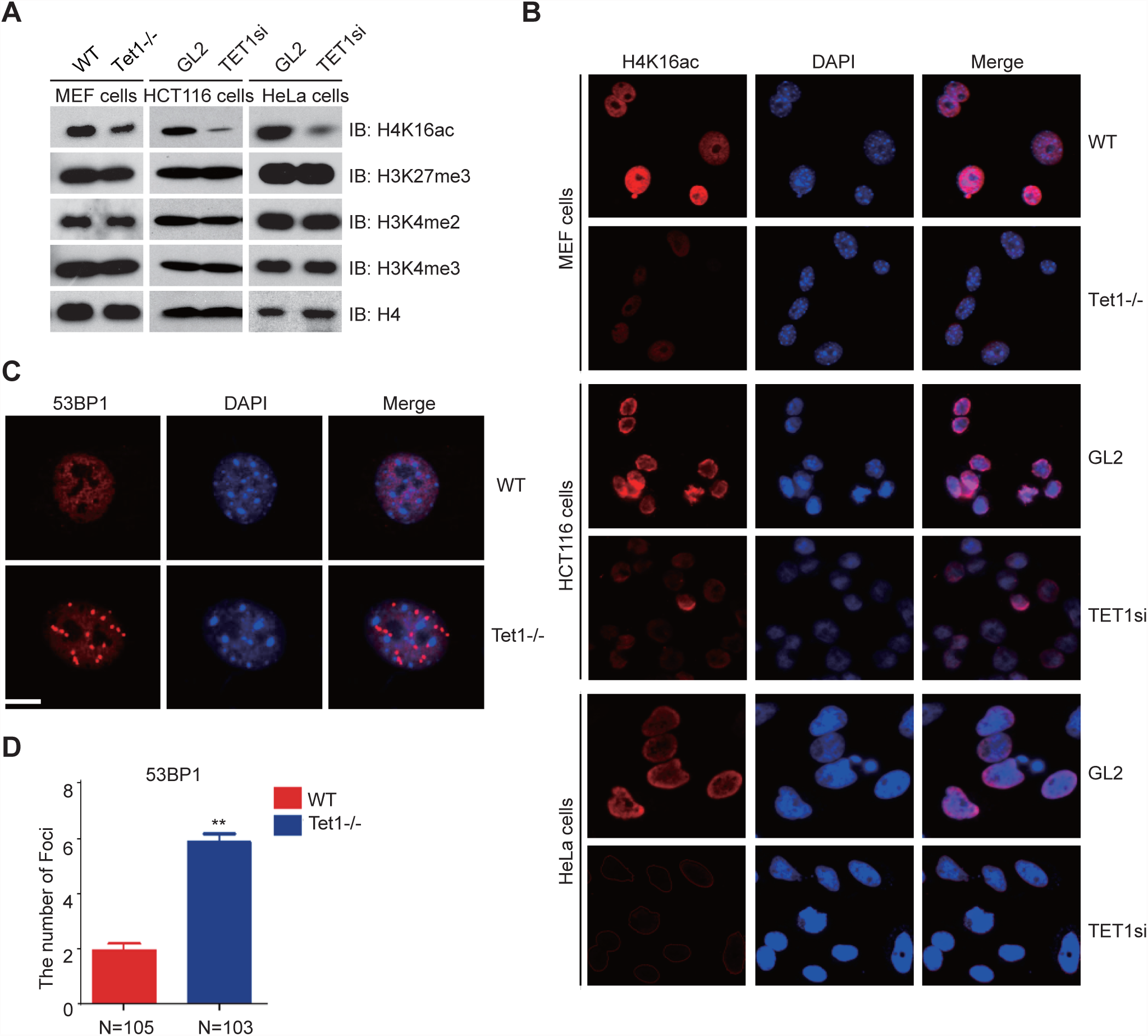
TET1 deficiency results in a reduction of H4K16ac levels. (**a**) Western blot analysis of histone modifications in Tet1^-/-^ MEF cells, TET1-knockdown HCT116 cells and TET1-knockdown HeLa cells using specific antibodies as indicated. (**b**) Immunofluorescence staining of Tet1^-/-^ MEF cells, TET1 knockdown HCT116 cells and TET1-knockdown HeLa cells with H4K16ac antibody. Nuclei were stained with DAPI. (**c**) Immunofluorescence staining of 53BP1 foci formation in Tet1^-/-^ MEF cells. Nuclei were stained with DAPI. (**d**) Statistical analysis of 53BP1 foci in Tet1^-/-^ MEF cells and WT MEF cells.

### TET1 forms a chromatin complex with SIN3A and hMOF to target H4K16ac

To confirm that TET1 formed a chromatin complex with hMOF and SIN3A, we first obtained different fractions of nuclear protein extracts in HEK293T cells separated by size fractionation using sucrose gradient centrifugation. As shown in our data, TET1, hMOF, and SIN3A were simultaneously enriched in pool 3, 4, and 5, suggesting they may be complexed with each other (Fig. 2a). Next, we performed immunoprecipitations (IPs) analysis of chromatin-bound protein after overexpression of Flag-TET1 or HA-SIN3A in HEK293T cells. Our data showed that both Flag-TET1 and HA-SIN3A interacted with hMOF (Fig. 2b). In order to identify which region of TET1 interacted with hMOF and SIN3A, we constructed three fragment plasmids of TET1, described as Flag-FL1, Flag-FL2, and Flag-FL3, which respectively contained CXXC domain, Cysteine-rich domain and DSBH (double stranded β-helix) conserved domain (Fig. 2c). Our Co-IP data showed that hMOF and SIN3A only interacted with Flag-FL3 (Fig. 2d). Consistently, the interactions were confirmed by His-pulldown assays using proteins overexpressed in, and purified from *E. coli* cells (Fig. 2e). Taken together, our results suggest that TET1 forms a chromatin complex with hMOF and SIN3A *via* its C-terminal region.

To determine whether TET1 targeted H4K16ac, we performed IP assays in HEK293T cells co-transfected with Flag-FL and HA-H4K16WT, HA-H4K16Q (a mimic acetylated mutant), or HA-H4K16R (an unacetylated mutant), respectively. Compared with interaction with HA-H4K16WT, Flag-FL had increased interaction with HA-H4K16Q, but decreased association with HA-H4K16R (Fig. 2f). Next, we performed chromatin fractionation and Western blot analysis on chromatin extracts from HEK293T cells with overexpressed HA-H4K16R and HA-H4K16WT, respectively. TET1 binding was significantly decreased in cells overexpressing HA-H4K16R compared with that in cells overexpressing HA-H4K16WT (Fig. 2g). In addition, when HEK293T cells were treated with the HDAC class I and class II inhibitor trichostatin A (TSA), increased TET1 binding was observed (Fig. 2h). Taken together, our data indicates that TET1 preferentially associates with histone H4 bearing K16 acetylation mark.

### TET1 depletion causes a significant reduction in H4K16ac levels

Given that TET1 contains CXXC domain, which enable its direct DNA binding, we reason that TET1 may recruit hMOF to genomic loci for regulation of H4K16ac mark. To test this hypothesis, we analyzed alterations of several histone modifications in TET1-depleted cells. Western blot analysis showed that H4K16 was hypo-acetylated in Tet1-knockout mice embryonic fibroblast (Tet1^-/-^ MEF) cells compared to in wild type Tet1^+/+^ MEFs. We observed a similar hypo-acetylation status in TET1-knockdown HCT116 and HeLa cells, respectively. However, the other histone markers, such as H3K4me2, H3K4me3, and H3K27me3, demonstrated insignificant alterations (Fig. 3a). Immunofluorescence staining further confirmed a significant reduction of H4K16ac in Tet1^-/-^ MEF cells, TET1-knockdown HCT116, and TET1-knockdown HeLa cells compared to their respective controls. Our results indicated that depletion of TET1 resulted in a significant reduction of H4K16ac level (Fig. 3b). Agree with this observation, the level of chromatin bound hMOF was significantly decreased in TET1-knockdown HEK293T cells (Fig. 4a).

**Figure 4.**
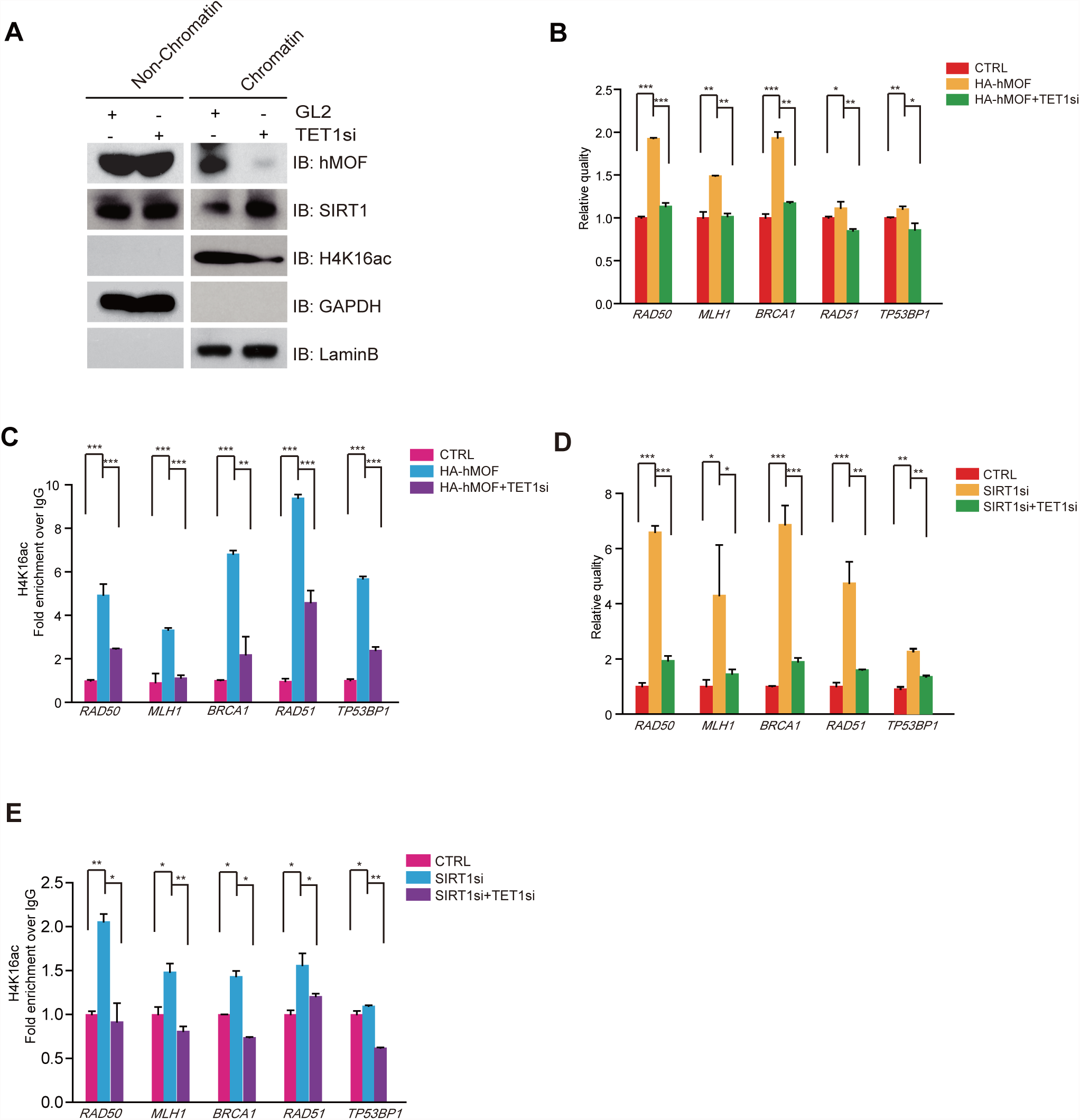
TET1 switches hMOF and/or SIRT1 binding on chromatin and regulates H4K16ac binding on the promoters of DNA repair genes. (**a**) Western blot analysis of the fractions in TET1-knockdown HEK293T cells using antibodies as indicated. (**b**) RT-qPCR analysis of *TP53BP1*, *RAD51*, *RAD50,* and *BRCA1* mRNA levels in HA-hMOF overexpressing HEK293T cells, with or without TET1-knockdown, as indicated (\**p*<0.05, \*\**p*<0.01, \*\*\**p*<0.001). (**c**) ChIP-qPCR analysis of H4K16ac binding at the promoter of *RAD50*, *BRCA1*, *RAD51*, and *TP53BP1* gene loci in HA-hMOF-overexpressing HEK293T cells, with or without TET1-knockdown, as indicated (\**p*<0.05, \*\**p*<0.01, \*\*\**p*<0.001). (**d**) RT-qPCR analysis of *RAD50*, *BRCA1*, *RAD51*, and *TP53BP1* mRNA levels in SIRT1-knockdown HEK293T cells, with or without TET1-knockdown, as indicated (\**p*<0.05, \*\**p*<0.01, \*\*\**p*<0.001). (**e**) ChIP-qPCR analysis of H4K16ac binding at the promoter of *RAD50*, *BRCA1*, *RAD51*, and *TP53BP1* in SIRT1-knockdown HEK293T cells, with or without TET1-knockdown, as indicated (\**p*<0.05, \*\**p*<0.01, \*\*\**p*<0.001). *p* values were calculated by unpaired Student’s *t* test.

As hypo-acetylation of H4K16 was reported to facilitate 53BP1 recruitment(Hsiao and Mizzen), we also determined the number of 53BP1 foci and found a two-fold increase in the number of 53BP1 foci in Tet1^-/-^ MEF cells (Fig. 3c and **3d**). Our results indicate that depletion of TET1 results in a significant reduction of H4K16ac levels and promotion the recruitment of 53BP1 binding to chromatin.

### TET1 switches binding of hMOF and/or SIRT1 on chromatin to regulate H4K16ac mark on the promoters of the DNA repair genes

Interestingly, using chromatin fractionation we found SIRT1 binding increased in TET1-knockdown HEK293T cells, whereas hMOF binding was decreased (Fig. 4a), indicating that TET1, as a switch, controls bindings of hMOF and SIRT1 to chromatin. To determine whether TET1 was required for regulation of DNA repair genes through modifying H4K16ac, we measured the mRNA levels of *RAD50, BRCA1, RAD51, and TP53BP1* in HA-hMOF overexpression or SIRT1-knockdown HEK393T cells, with or without TET1 knockdown, *via* RT-qPCR. We found that depletion of TET1 blocked the increased expression of *RAD50*, *BRCA1*, *RAD51*, and *TP53BP1* both in hMOF-overexpressing cells (Fig. 4b) and in SIRT1-knockdown cells (Fig. 4d). Next, we further determined the effect of H4K16ac enrichment at the promoter of *RAD50*, *BRCA1*, *RAD51*, and *TP53BP1* in HA-hMOF-overexpressing or SIRT1-knockdown HEK293T cells, with or without TET1 knockdown, *via* ChIP-qPCR. The results showed that depletion of TET1 blocked the increased H4K16ac enriched at the promoter of *RAD50*, *BRCA1*, *RAD51*, and *TP53BP1* in both hMOF-overexpressing (Fig. 4c) and SIRT1-knockdown HEK293T cells (Fig. 4e). These results suggest that TET1, as a switch, inversely modulates the bindings between hMOF and SIRT1 on chromatin, which in turn dynamically controls H4K16ac levels, and ultimately contributes to the expression of these important DNA repair genes.

### Impairment of homologous recombination repair and non-homologous end joining in TET1-, but not TET2- or TET3-knockdown cells

To determine the role of TET1 in DNA repair, C57 wild type mice (WT) and Tet1 heterozygous mutated mice (Tet1^+/-^ mice) were subjected to x-ray radiation. The coat-state rating scale results showed that there was a severe deterioration of the coat in Tet1^+/-^ mice compared with WT mice after four months of x-ray radiation (Fig. 5a, 5b), suggesting that Tet1^+/-^ mice had defects in DNA repair mechanisms in response to DSBs. Next, we measured the percentage of DNA present in comet tail and the tail moment in comet assay to determine the extent of DNA damage in Tet1^-/-^ MEF cells, respectively. Our results showed both of these parameters were increased approximately by two-folds in the Tet1^-/-^ MEF cells than those in WT cells (Fig. 5c, 5d, and 5e), indicating that Tet1^-/-^ MEF cells had a higher level of DSBs. We also measured foci formation of DSBs maker γH2AX by immunofluorescence staining. Consistent with our comet assay, we found that the number of γH2AX foci increased two-fold in Tet1^-/-^ MEF cells compared to WT cells (**Supplementary Fig4a, 4b**). In addition, Western blot analysis indicated that the level of γH2AX increased in TET1-knockdown cells, but not in TET2- or TET3-knockdown HEK293T cells (**Supplementary Fig.4c**). DAPI staining and statistical analysis further showed that the percentage of micronuclei in Tet1^-/-^ MEF cells were two-folds higher than that in WT cells (Fig. 5f, 5g). These results indicate that loss of TET1 leads to severe DNA damage and the defects in DNA repair and genomic instability.

**Figure 5.**
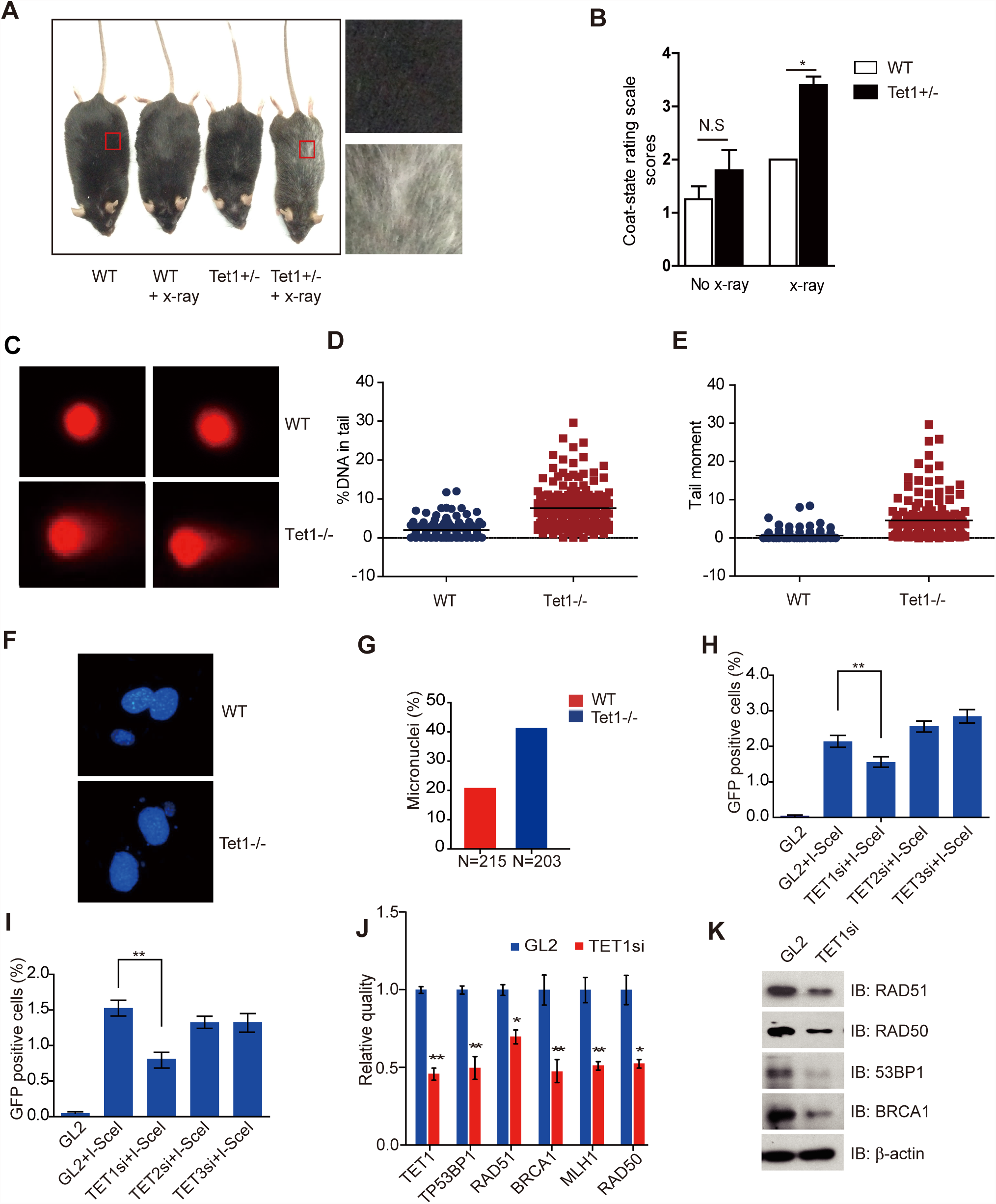
Depletion of TET1 impairs HRR and NHEJ and downregulation of DNA repair genes. (**a**) The phenotype of WT mice and Tet1^+/-^ mice after 3Gy x-ray irradiation. Mice were irradiated with a single whole-body dose of 3Gy x-rays at 60 days of age. Concurrent sham-irradiated control groups were also examined from the same litter where possible to minimize genetic bias. (**b**) Statistical analysis of the coat-state condition of mice after 3Gy x-ray radiation, according to the method published by C Nasca, *et al*. (no x-ray: WT n=14, Tet1^+/-^ n=15; x-ray: WT, n=13, Tet1^+/-^, n=10; \**p*<0.05, N.S *p*>0.05). (**c**) DNA damage in WT and Tet1^-/-^ MEF cells, as measured by neutral comet assay. (**d, e**) Quantification of comet experiments (shown in **a**) the percentage of DNA in the comet tail or the tail moment was measured and statistical analysis was performed using GraphPad software. (**f**) DAPI staining and microscope analysis of micronuclei in Tet1^-/-^ MEF cells. (**g**) Statistical analysis of micronuclei in WT and Tet1^-/-^ MEF cells (\*\**p*<0.01, N = the number of the cells). (**h**) Frequency of HRR after TET1, TET2, or TET3 depletion (\*\**p*<0.01). (**i**) Frequency of NHEJ after TET1, TET2, or TET3 depletion (\*\**p*<0.01). (**j**) RT-qPCR analysis of mRNA levels in DNA repair genes (*RAD50*, *BRCA1*, *RAD51*, and *TP53BP1*) in TET1-knockdown cells (\**p*<0.05, \*\**p*<0.01). (**k**) Western blot analysis of 53BP1, RAD51, RAD50, and BRCA1 protein expression in TET1-knockdown HEK293T cells. *p* value was calculated by unpaired Student’s *t* test.

Homologous recombination repair and non-homologous end joining are two types of DNA repair mechanism in response to DSBs. To determine the extent of HRR and NHEJ repair frequencies in TET1-depleted cells, we set out to employ two types of GFP reporter systems in HEK293T cells, in which a defective GFP gene is functionally restored to WT cells upon I-SceI transfection(Mao et al. 2008). We found TET1-depletion resulted in 25% decrease of GFP positive cell numbers in HRR reporter assay (Fig. 5h), and 50% reduction in NHEJ reporter assay, and neither of these outcomes was observed in TET2- and TET3-knockdown cells (Fig. 5i). More importantly, we demonstrated that TET1-depletion induced transcriptional repression of the important genes in DNA repair, including *RAD50*, *BRCA1*, *RAD51*, and *TP53BP1*, *via* RT-PCR and Western blot analysis (Fig. 5j, 5k). Collectively, our results indicate that depletion of TET1, but not TET2 or TET3, results in the defects of HRR and NHEJ in response to DSBs and genomic instability through downregulation of the DNA repair genes.

### Dynamic regulation of TET1, SIN3A, hMOF, and SIRT1 binding on chromatin in response to hydrogen peroxide-induced oxidative damage

Next, we investigated alterations in different histone modifications by treating HEK293T cells with DNA damaging reagents including adriamycin (ADR), bleomycin (Bleo), camptothecin (CPT), and hydrogen peroxide (H_2_O_2_). Our data showed that H_2_O_2_ treatment specifically resulted in a significant decrease of H4K16ac levels, whereas H3K27me3, H3K4me2, and H3K4me3 had no obvious changes in response to DNA damage reagents (Fig. 6a). In addition, we measured the bindings of TET1, hMOF1, SIRT1, and SIN3A on chromatin after DNA damage reagent treatment as described above. We found that H_2_O_2_ treatment caused a significant decrease of both TET1 and hMOF binding, and an increase in SIRT1 binding, but no obvious alteration of SIN3A binding (Fig. 6b). To support this notion, H_2_O_2_ treatment caused significantly decreased interactions between Flag-FL3 with hMOF, but not with SIN3A (Fig. 6c). As shown in Fig. 6d, H_2_O_2_ treatment also led to transcriptional repression of the important genes in DNA repair including *RAD50*, *BRCA1*, *RAD51*, and *TP53BP1*, while the DNA methylation level, as assessed by profiling using reduce representation bisulfite sequencing (RRBS), did not present alteration in their corresponding promoters (Fig. 6e, 6f). ChIP-qPCR assay further revealed that TET1 and H4K16ac enriched together at promoter regions of these DNA repair genes and displayed decreased enrichment after H_2_O_2_ treatment (Fig. 6g). Taken together, our results suggest that oxidative damage induces dynamic recomposition of TET1, hMOF, and SIRT1 binding on chromatin, which is concurrent with the deacetylation of H4K16 and reduced expression of the important DNA repair genes.

**Figure 6.**
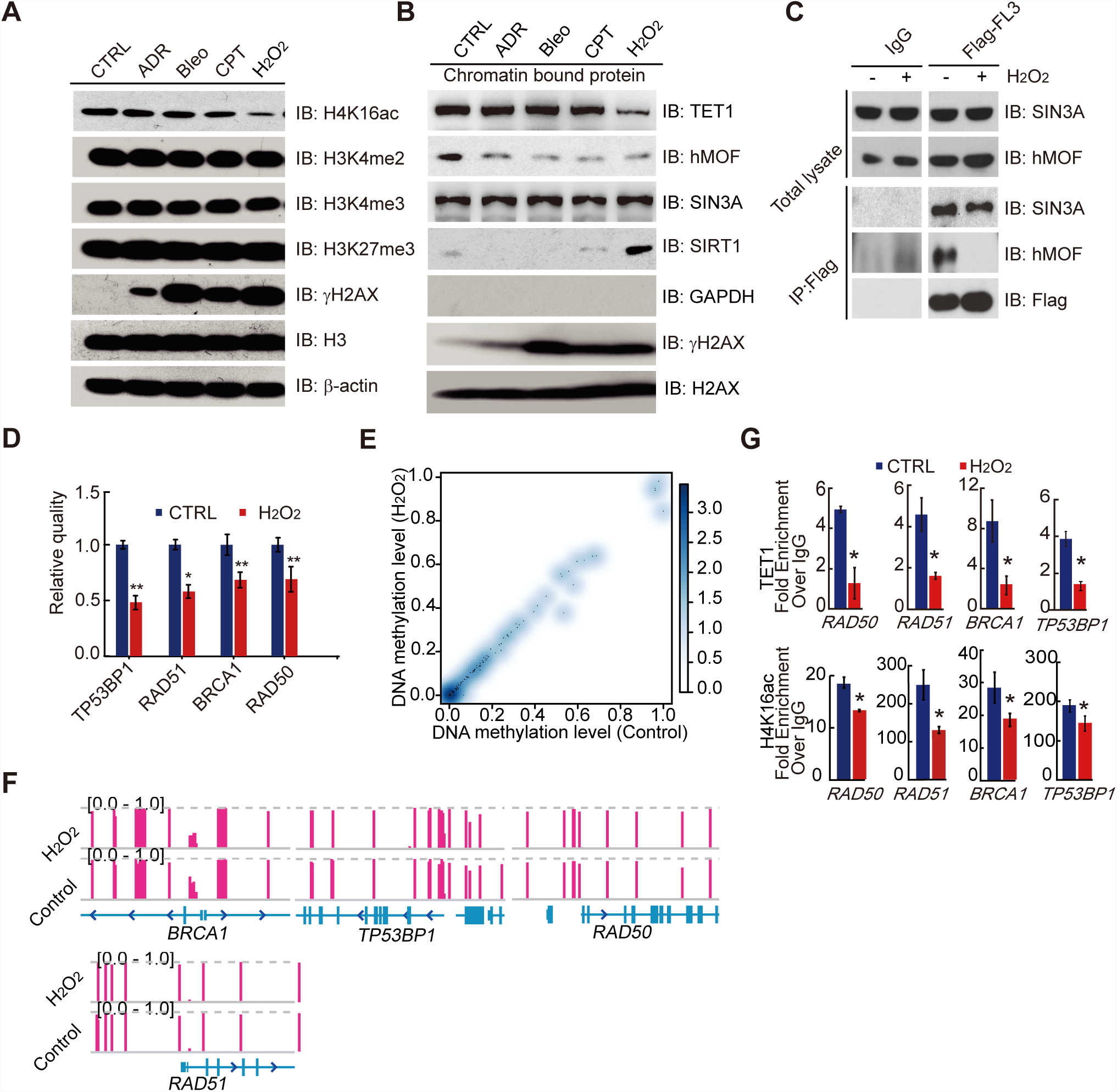
Dynamic regulation of TET1, SIN3A, hMOF, and SIRT1 binding on chromatin in response to H_2_O_2_-induced oxidative damage. (**a**)Western blot analysis of histone modifications levels, using specific antibodies, after 1μM ADR, 1μM Bleo, 1μM CPT, and 2mM H_2_O_2_ treatments for 1h, respectively. (**b**)Western blot analysis on chromatin fraction using antibodies as indicated after ADR, Bleo, CPT, and H_2_O_2_ treatments for 1h, respectively. γH2AXand H2AX were used as a DNA damage marker and a chromatin fraction marker, respectively. GAPDH was used as a cytoplasm fraction marker. (**c**) HEK293T cells were transfected with Flag-FL3 and then untreated or treated with H_2_O_2_. IPs were performed using a Flag-tag antibody, followed by Western blot analysis using the indicated antibodies. (**d**) RT-qPCR analysis of mRNA levels in *RAD50*, *BRCA1*, *RAD51*, and *TP53BP1* in untreated and H_2_O_2_-treated HEK293T cells (\**p*<0.05, \*\**p*<0.01). (**e**) Smooth scatterplot of DNA methylation levels at the promoter of DNA repair genes in HEK293T cells, with or without H_2_O_2_ treatment, through RRBS. Pearson Correlation Coefficient =1. (**f**) IGV map of the DNA methylation levels at the TSS of *RAD50*, *BRCA1*, *RAD51*, and *TP53BP1*. Both the methylation level at promoters and flanking regions were displayed on two IGV tracks, one for the control group and the other for H_2_O_2_ treatment group in HEK293T cells. (**g**) ChIP-qPCR verification of TET1 and H4K16ac binding before and after H_2_O_2_ treatment in HEK293T cells. *p* value was calculated by unpaired Student’s *t* test.

### TET1 binding on chromatin is dependent on ataxia telangiectasia mutated (ATM) protein in HEK293T cells

Our data also showed an increased TET1 binding on chromatin after wortmannin treatment, which is an inhibitor of early DNA damage response proteins PIKK family (phosphatidylinositol 3-kinase–related protein kinase) (**Supplementary Fig. 5a**), indicating that TET1 binding was dependent on PIKK family kinases. Further studies showed that there was increased binding of TET1 after ATM inhibition using either its inhibitor CGK733 or ATM-siRNA (**Supplementary Fig. 5b, 5c**), suggesting that binding of TET1 on chromatin is dependent on ATM, but not DNA-PKcs (**Supplementary Fig. 5d**).

### A similar epigenetic TET1/SIN3A/hMOF recomposition mechanism is verified in an H-RAS^V12^ transformed cell line

Previous studies have reported that oxidative damage leads to a higher risk of cancers in humans by diminishing histone acetylation, which predominantly occur at H4K16ac(Leufkens et al.; Fraga et al. 2005; Sosa et al. 2013). Therefore, we hypothesized that dynamic recomposition of this TET1/SIN3A/hMOF/SIRT1 chromatin components might contribute to tumorigenesis. By employing an epithelial ovarian cell line T29 and its oncogenic counterpart T29H cell line which is transformed with H-RAS^V12^, we simultaneously analyzed the levels of oxidation and H4K16ac, expression of DNA repair genes, and the binding of TET1, SIN3A, hMOF1, and SIRT1, as compared with those in WT cells. Consistently, transcriptional expression of *RAD50*, *BRCA1*, *RAD51*, and *TP53BP1* was decreased in T29H cells (Fig. 7a), but the methylation levels on their promoters was not significantly altered according to our RRBS data (**Supplementary Fig. 6a**). As expected, compared to T29 cells, there were decreased bindings of TET1 and hMOF, and increased SIRT1 binding on chromatin in T29H cells (Fig. 7b), concurrent with the reduction of H4K16ac levels (Fig. 7c). The bindings of both TET1 and H4K16ac on the promoter of these DNA repair genes were decreased as well (Fig. 7d, 7e). Meanwhile, we also found elevated 8-OH-dG level using a dot blot assay in T29H cell (**Supplementary Fig. 6b**). These results suggested that oncogenic transformation causes hypoacetylation of H4K16 *via* decreased chromatin binding of TET1 and hMOF1, and increased binding of SIRT1, which lead to downregulation of the DNA repair genes, ultimately impairing DNA repair and involving tumorigenesis.

**Figure 7.**
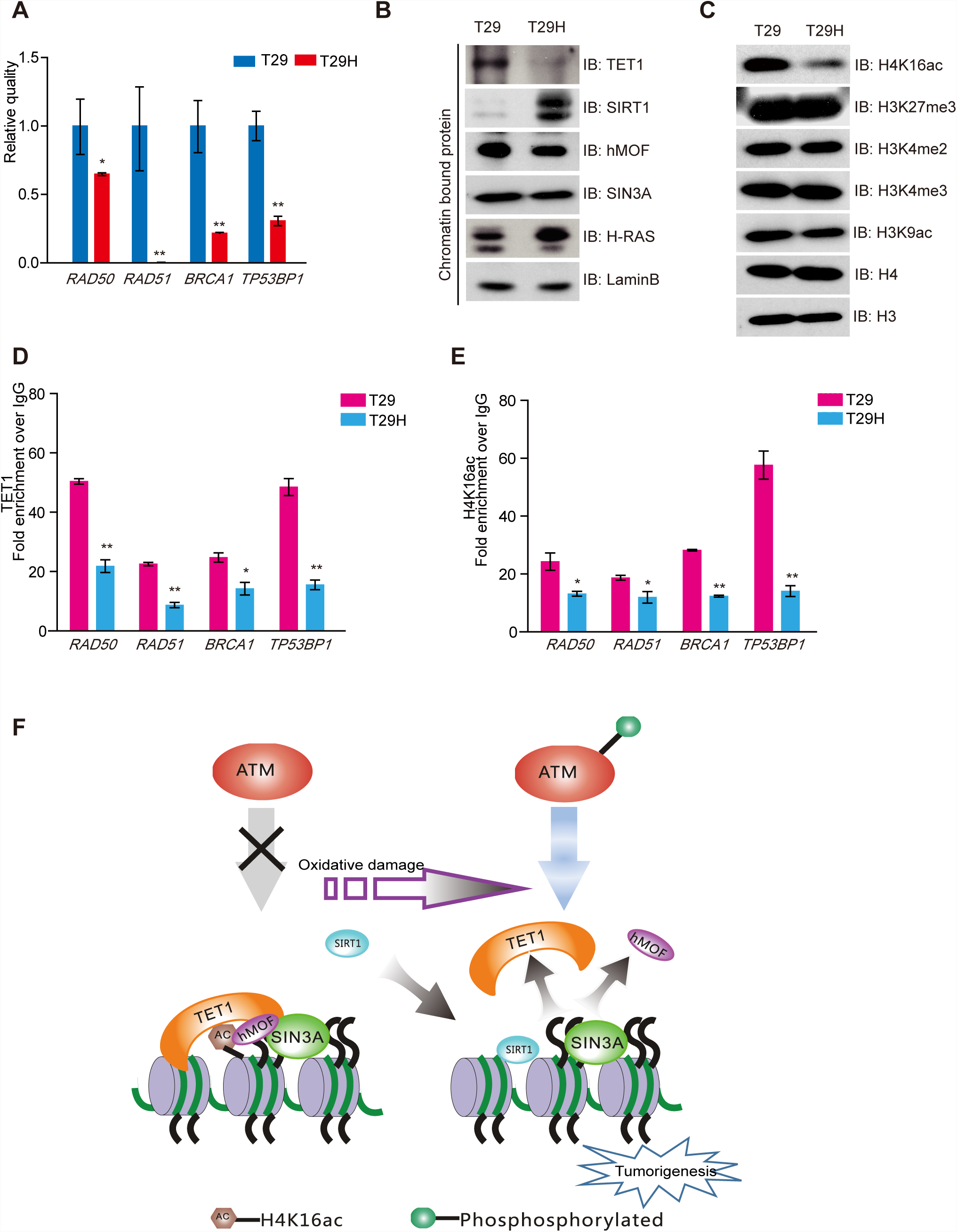
Similar recomposition mechanism of the TET1/SIN3A/hMOF complex was observed in an oncogenic transformed cell line. (**a**) RT-qPCR analysis of mRNA levels of *53BP1*, *RAD51*, *RAD50*, and *BRCA1* in T29 and T29H cells, respectively (\**p*<0.05, \*\**p*<0.01). (**b**) Western blot analysis of TET1, SIRT1, hMOF, and SIN3A bindings on chromatin in T29 and T29H cells, respectively. (**c**) Western blot analysis of histone modifications in T29 and T29H cells using specific antibodies as indicated. (**d, e**) ChIP-qPCR analysis of both TET1 and H4K16ac binding on the promoter of *RAD50, RAD51*, *BRCA1*, and *TP53BP1* in T29 and T29H cells (\**p*<0.05, \*\**p*<0.01). (**f**) Working model of the dynamic regulation of TET1, SIN3A, hMOF, and SIRT1 *via* targeting H4K16ac in response to oxidative damage.

## DISCUSSION

In this study, we revealed that TET1 could function as a core component of TET1/SIN3A/hMOF chromatin complex, supported by their co-occupancy of common targets across the genome, association with each other by co-IP, and co-migration in size fractionation assays (Fig. 1, 2a-e). However, we still cannot rule out the possibility that TET1, hMOF, and SIN3A form sub-complexes to co-occupy the same genomic regions. We demonstrated that TET1 controlled the bindings of hMOF and SIRT1 on chromatin to specifically modulate alteration of H4K16ac. H_2_O_2_-induced oxidative damage could induce recomposition of this chromatin complex to provoke hypoacetylation of H4K16, which suppressed the expression of the important DNA repair genes and ultimately impaired HRR and NHEJ. Consistently, by employing H-RAS^V12^ oncogenic transformed cell line, we revealed a similar epigenetic recomposition mechanism for understanding the role of TET1 in tumorigenesis. With this work, a novel role of TET1 is proposed as shown in Fig. 7f.

Previous reports showed TET1 and modified 5-hmC correlates with “bivalent” chromatin markers with both repressing (H3K27me3) and activating (H3K4me3) chromatin markers in mESCs(Pastor et al. 2011; Wu et al. 2011). The fact that TET1 contributes to silencing some genes by facilitating recruitment of PRC2 complex(Wu et al. 2011), supports the hypothesis the TET1 represses gene expression by forming a TET1/PRC2 repressor complex that targets H3K27me3. However, the following two lines of evidence imply that involvement of TET1/PRC2 in transcriptional repression may be ESCs-specific: 1) correlation between 5hmC and H3K27me3 is unique to ESCs, and is not present in differentiated fibroblasts or adult tissues; and 2) interaction between TET1 and the components of PRC2 complex were only observed in ESCs, but not in fibroblasts or HEK293T cells(Neri et al. 2013). Our observation, in which co-occupancy between Tet1 and Suz12/Ezh2/H3K27me3 present in cluster M7 (mainly involved in cell differentiation and cell fate functions) (**Supplementary Fig. 2c**), further supports the notion that Tet1/PRC2 complex may play an important role in cell differentiation and cell fate in mESCs. Additionally, we observed high co-occupancy of Tet1/Mof/Sin3a/H4K16ac at DNA repair genes in cluster M6, whereas binding of H3K27me3 was absent (**Fig.1**). Our results, combined with a previous report in which TET1 was shown to interact with SIN3A in HEK293 cells and their shared binding on TSS region are H3K27me3-binding negative(Neri et al. 2013), exclude the possibility that TET1 downregulates these DNA repair genes by TET1/PRC2 complex through activating the repressive histone mark H3K27me3 in differentiated cells.

Recent studies demonstrate that TET1 acts as a stable component of O-GlcNAc transferase (OGT) in ESCs, and promotes histone O-GlcNAcylation and H3K4me3 states for gene activation(Vella et al. 2013). However, several reports argue that TET2 and TET3, but not TET1, interact with OGT to activate gene expression in HEK293T cells(Chen et al. 2005; Deplus et al. 2013), raising the possibility that TET1/OGT complex might be involved in transcriptional activation exclusively in ESCs. Our results in several differentiated cells lines revealed that TET1 knockdown resulted in hypoacetylation of H4K16 only, but did not affect levels of H3K27me3, H3K4me2, and H3K4me3, suggesting that TET1 complex mainly modulate H4K16ac in these cells (**Fig.3**). Interestingly, the observations that inhibition of HDAC1/HDAC2 has no significant effect on some important genes in DNA repair, including *RAD50* and *BRCA1*(Thurn et al. 2013), and HDAC1/HDAC2 do not have occupancy on the promoter of *RAD50*, *BRCA1*, *RAD51*, and *TP53BP1* in ChIP-seq data from mESCs (**Supplementary Table 2**), imply that TET1/SIN3A may not form a complex with HDAC1/HDAC2 to repress these DNA repair genes.

One of main etiological hypothesis linking genomic instability, mutagenesis and tumorigenesis is that deficient DNA repair mechanisms is derived from extensive oxidative DNA damage and cellular injury(Ziech et al. 2011; Gencer et al. 2012). The relationship between DNA damage and epigenetic gene silencing has been examined using an engineered cell model, in which an I-SceI restriction site was integrated into the CpG island of the E-cadherin promoter(O’Hagan et al. 2008). That study showed that H_2_O_2_-induced oxidative damage regulated epigenetic DNA methylation changes through redistribution of DNMT1, EZH2, and SIRT1 on chromatin from non-CpG rich regions to CpG islands(O’Hagan et al. 2011). Here, we observed that H_2_O_2_-induced oxidative damage directly resulted in 165 differentially methylated regions (DMRs) in promoters (data not shown); however, none of these involved in genes in the DNA repair pathway (Fig. 6). These results further support that downregulation of expression of *RAD50*, *BRCA1*, *RAD51*, and *TP53BP1* under H_2_O_2_-induced oxidative damage, is independent of promoter methylation changes on those genes. Interestingly, we show that SIRT1 binding is elevated under both H_2_O_2_-induced oxidative damage and oncogenic transformation, and SIRT1 forms a chromatin complex with PRC2 complex subunits in response to H_2_O_2_-induced oxidative damage, which is consistent with a previous report(O’Hagan et al. 2011) (**Supplementary Fig. 7a**). This supports the hypothesis that epigenetic silencing of important genes in DNA repair both in cancer cells and under oxidative damage, likely results from loss of TET1 which switches the bindings of hMOF and SIRT1 on chromatin, thus promoting aberrant hypoacetylation of H4K16, rather than the alteration of DNA methylation. This hypothesis is further supported by the observation that a similar epigenetic dynamic alteration occurred in H-RAS^V12^oncogenic transformed cells (Fig. 7). Stephen P. Jackson’s group showed that H3K4me3, H3K18ac, H4K5ac, and H4K12ac were unaffected in response to oxidative damage(Tjeertes et al. 2009). Our results also suggested that it’s unlikely that hypoacetylation of H3K9ac plays a repressive role in regulation of important genes in DNA repair in response to H_2_O_2_-induced oxidative damage (**Supplementary Fig. 6b**). These evidence suggest that hypoacetylation of H4K16 is likely a major mechanism to downregulate these DNA repair genes under oxidative damage. However, the repressive role of other histone marks and microRNAs cannot be ruled out and need to be further explored.

## MATERIALS and METHODS

### Mice

Tet1^+/-^ mice are obtained from the Jackson Laboratory (Cat# 017358). For genotyping of Tet1^+/-^ mice, the forward primer AACTGATTCCCTTCGTGCAG, and the reverse primer TTAAAGCATGGGTGGGAGTC were used. The expected band size for homozygote mutant was 650bp, 850bp for the wild type strain, and 650bp and 850bp double bands for the heterozygote strain.

### X-ray irradiation

Irradiations were performed at the Chinese Academy of Sciences (Beijing)using an x-ray machine (RS-2000 PRO Biological system). WT mice and Tet1^+/-^ mice were irradiated with a single whole-body dose of 3Gy x-ray at 60 days of age. The irradiation was operating at 160-kV constant potential and 25 mA (0.3 mm Cu filter) at a dose rate equal to 1.136Gy min–1 for a total of 2.64 min. The cage was cleaned with 75% ethanol when each of irradiation was finished. Coat–state condition of WT mice and Tet1^+/-^ mice with irradiation or sham-irradiated were scored after four month of irradiation.

### Cell Culture, Plasmids, and siRNA Oligonucleotides

Mouse Tet1^-/-^ MEF cells were a generous gift from Dr. Guoliang Xu, SIBS of Chinese Academy of Sciences, Shanghai. T29 and T29H cells were a generous gift from Dr. Jason Liu from MD Anderson Cancer Center of University of Texas, Houston. All the cells were cultured in DMEM media (Hyclone, USA), supplemented with 10% fetal bovine serum (FBS; Hyclone, USA) and penicillin-streptomycin (Invitrogen, USA). H4K16Q and H4K16R mutants were generated using a QuickChange Multi Site-Directed Mutagenesis Kit (Stratagene, USA). RNA interference (RNAi) experiments were performed using Dharmacon siGENOME SMARTpool siRNA duplexes (Thermo Fisher Scientific, USA) against *TET1*, *TET2*, *TET3*, *HDAC1*, *HDAC2*, *ATM*, and *SIRT1* respectively. *TET1* cDNA was purchased from Origene (RC218608), cDNA for *hMOF, SIRT1, SIN3A*, and *Histone H4* were generated from cDNA library and subcloned into pcDNA3.0 vector, followed by sequencing validation. Antibodies used in this study were purchased from different commercial companies as detailed in Supplemental Methods.

### siRNA Transfection, RNA Isolation, and qRT-PCR

See Supplemental Methods for detailed information.

### ChIP-seq Data Preparation

For integrated genomic analysis, we collected 219 ChIP-seq data sets of 103 DNA binding proteins and 14 data sets of 8 histone modification markers in mESCs from GEO and ENCODE (See **Supplementary Table 1**). Further analysis details are provided in Supplemental Methods.

### RRBS Library Preparation, Sequencing and Analysis

See Supplemental Methods for detailed information.

### ASSESSION

GSE66395

## AUTHOR CONTRIBUTIONS

J.N.Z. and Z.S.S. designed the experiments; S.S.D. and J.A.Z. performed immunofluorescence and microscopic analysis; K.L.W. helped W.S.C. with ChIP-Seq and RRBS; X.F.L. and F.B.M. performed ChIP–Seq and RRBS data analysis; S.S.D. and J.A.Z. helped for immunoprecipitation and cellular fractionation experiments. J.Y. helped for gene cloning. Z.C, Y.Y.L. and Y.W. helped for mice experiments. J.N.Z. and Z.S.S. wrote the manuscript; H.M.Y., X.Z.X. and J.Y.W. revised the manuscript.

## ACKNOWLEDGMENTS

We are grateful to Dr. Guoliang Xu for a kind gift of TET1 knockout MEF cells, Dr. Jie Du for critically evaluating the manuscript, and Dr. Xingda Ju and the other members in our lab for help in experiments. This work is supported by National High Technology Research and Development Program of China (No. 2012AA02A202); Natural Science Foundation of China (NSFC)-Canadian Institutes of Health Research Collaborative Research Project (No. 81161120541); Nation Natural Science Foundation of China (NSFC)(No. 81301778); Zhejiang Provincial Natural Science Foundation (ZNSF) (No. LY13C060002); Zhejiang Provincial Department of Education Research Project (No. 84612051); Nation Natural Science Foundation of China (NSFC) (No. 31271266); Startup Fund for scholars of Wenzhou medical university (No. QTJ12001), and grant from Innovative Center of China, AstraZeneca.

## SUPPLEMENTARY MATERIAL

**Supplementary Figure 1. Enrichment analysis of Tet1, Sin3a, Mof, and H4K16ac** Heat-map showed a ChIP-seq signal enrichment visualization of Tet1, Sin3a, Mof, and H4K16ac in mESCs.

**Supplementary Figure 2. Binding states of Tet1/Sin3a/Mof, PRC2/Suz12/Ezh2, and Sin3a/Hdac1/Hdac2 complexes and its function enrichment analysis in the promoter regions** (**a**) The seven binding states of TET1 as ordered by emission and transition parameters through ChromHMM software. (**b**) A Chow-Ruskey diagram of the overlapped genes of 5 binding states (M2, M3, M5, M6, and M7). (**c**) Top 60 biological processes and KEGG pathways of the 5 binding states. *p* values were converted to -10*log (*p* value).

**Supplementary Figure 3. ChIP-seq signals of Tet1, Sin3a, Mof, and H4K16ac in promoter regions of the DNA repair genes** IGV map presented ChIP-seq signals of Tet1, Sin3a, Mof, and H4K16ac in promoter regions of some DNA repair genes, including *Brca1*, *Brca2*, *Rad50*, *Rad51*, and *Mlh1*.

**Supplementary Figure 4. Increased γH2AX in TET1-depleted cells** (**a**) DNA damage marker γH2AX was stained in WT and Tet1^-/-^ MEF cells. Nuclei were stained with DAPI. (**b**) Statistical analysis of γH2AX foci pictured in (**a**). More than ten number of foci was calculated as ten (\*\**p*<0.01, N = the number of the cells). (**c**) Western blot analysis of γH2AX level in TET1-, TET2-, or TET3-knockdown HEK293T cells.

**Supplementary Figure 5. Binding of TET1 on chromatin is dependent on ATM** (**a**) Western blot analysis of the fractions before and after 1μM wortmannin treatment in HEK293T cells. (**b**) Western blot analysis of the fractions before and after 1μM CGK733 in HEK293T cells. (**c)** Western blot analysis of the fractions of the control and ATM siRNA-knockdown using indicated antibodies in HEK293T cells. (**d**) Western blot analysis of the fractions before and after 1μM DNA-PKcs inhibitors NU7026 treatment using indicated antibodies in HEK293T cells.

**Supplementary Figure 6. Analysis of DNA methylation on the promoter of the DNA repair genes and oxidative damage levels in T29 and T29H cells** (**a**) DNA methylation levels of *RAD50*, *MLH1*, *BRCA1*, *RAD51*, and *TP53BP1* genes in T29 and T29H cells through RRBS. (**b**) Dot blot analysis of oxidative damage level in T29 and T29H cells using an 8-OH-dG antibody.

**Supplementary Figure 7. Interactions between SIRT1 and PRC2 complex and binding of H3K9ac on the promoter of the DNA repair genes before and after H_2_O_2_ treatment** (**a**) Nuclear protein IPs were performed using SIRT1 antibody with or without 2mM H_2_O_2_ treatment and then detected by Western blot analysis using SUZ12 and EZH2 antibodies. (**b**) ChIP-qPCR analysis of H3K9ac binding on the promoter of *RAD50, MLH1*, *RAD51*, *BRCA1*, and *TP53BP1* with or without H_2_O_2_ treatment in HEK293T cells (N.S *p*>0.05).

**Supplementary Table 1.** Summary of published ChIP-seq data sets of DNA-binding proteins and histone modifications in mESCs

**Supplementary Table 2.** Occupancies of Tet1, Mof, Sin3a, Hdac1, Hdac2, H4K16ac, and H3K27me3 on the promoter of DNA repair genes from ChIP-seq data sets in mESCs

